# A role for KIF9 in male fertility

**DOI:** 10.1101/2020.03.21.001602

**Authors:** Ken Chen, Sang Yeon Cho, Yongwei Zhang, Amanda Beck, Jeffrey E. Segall

## Abstract

A mouse was generated containing a floxed exon 3 of the gene for the kinesin family member KIF9. By in situ hybridization, expression of KIF9 mRNA was highest in the testis and was also strong in epithelia containing multi-ciliated cells such as the ependyma, bronchioles and oviduct. Deletion of the exon led to loss of KIF9 expression at the mRNA and protein level with no effect on viability. However, homozygous KIF9 knockout males were sterile. Although KIF9 knockout sperm were motile, they were unable to fertilize oocytes in an in vitro fertilization assay. Closer examination of sperm motility indicated a subtle difference in waveform. Our results suggest that KIF9 plays a role male fertility, possibly through regulation of flagellar waveforms in ciliated cells.

## Introduction

In mammalian cells, the kinesin family member 9, KIF9, has been reported to contribute to the ability of macrophages to degrade extracellular matrix[1], potentially through the regulation of delivery of matrix degrading enzymes to the cell surface. In addition, suppression of KIF9 expression has been reported to have effects on spindle dynamics[2]. Intriguingly, KIF9 is overexpressed in glioblastoma and correlates with poor outcome in that disease[3]. Based on our observations that microglia (a macrophage subtype found in the brain) could enhance glioma invasion[4], it is possible that KIF9 plays a role in the aggressiveness of cancer through enabling macrophages to stimulate tumor cell invasion.

Conversely, in lower eukaryotes, KIF9 homologues have been shown to play a role in ciliary motility. The first KIF9 homologue identified (KLP1), was found as a flagellar protein of Chlamydomonas reinhardtii[5]. Knockdown of KLP1 in Chlamydomonas results in a strong decrease in flagellar motility[6]. Similarly, loss of KIF9 homologues in other lower eukaryotes such as trypanosomes or fern gametocytes leads to reduced flagellar based motility.

To determine whether KIF9 plays an important role in cancer biology, we decided to generate a floxed KIF9 mouse that would allow us to study to potential roles of stromal KIF9 in the invasion of glioblastoma and other cancers. During the development of this mouse model, we found a strong male sterility defect consistent with a role in flagellar motility, and report on that defect here. A similar observation has recently been reported[7].

## Methods

### Generation of floxed KIF9 mice

All animal procedures were performed according to NIH guidelines and approved by the Committee on Animal Care at Einstein. A pair of gRNAs targeting intron 2 (gRNAE22) and intron 3 (gRNAE23) of KIF9 gene respectively were designed using an online tool (https://benchling.com/) and generated by in vitro transcription[1] (Table 1 and Figure 1). Cas9 protein was purchased from PNA Bio Inc. A KIF9 conditional knockout(CKO) HRD plasmid carrying around 2kb homologous arms on each side surrounding a floxed KIF9 exon3 with an added primer sequence (KS) for genotyping (Supplementary Data 1) was generated by SLiCE cloning[2]. Superovulated female C57BL6 mice (3–4 weeks old) were mated to C57BL6 males, and fertilized embryos were collected from oviducts. The gRNAs targeting introns 2 and intron 3 of the KIF9 gene, the Cas9 protein and the KIF9 conditional knockout(CKO) HRD were mixed and microinjected into the pronuclei of fertilized eggs. The injected zygotes were transferred into pseudopregnant CD1 females, and the resulting pups were genotyped. Founder mice carrying floxed KIF9 alleles were identified by pcr using primers 3’SR and 5’SF and sequencing. Founders were mated with C57BL6 mice to get F1 offspring. Crossing of F1 floxed heterozygotes was used to generate homozygous floxed mice. KIF9 knockout mice were generated by breeding floxed homozygotes with C57BL/6-Tg(Zp3-cre)93Knw/J mice (Jackson Labs catalogue # 003651). Genotyping was performed by PCR of genomic DNA using the 3’SR and 5’SF primers, with product sizes used to determine genotype (1003 bp for WT, 1085 bp for floxed, 593 for knockout).

**Table 1:**
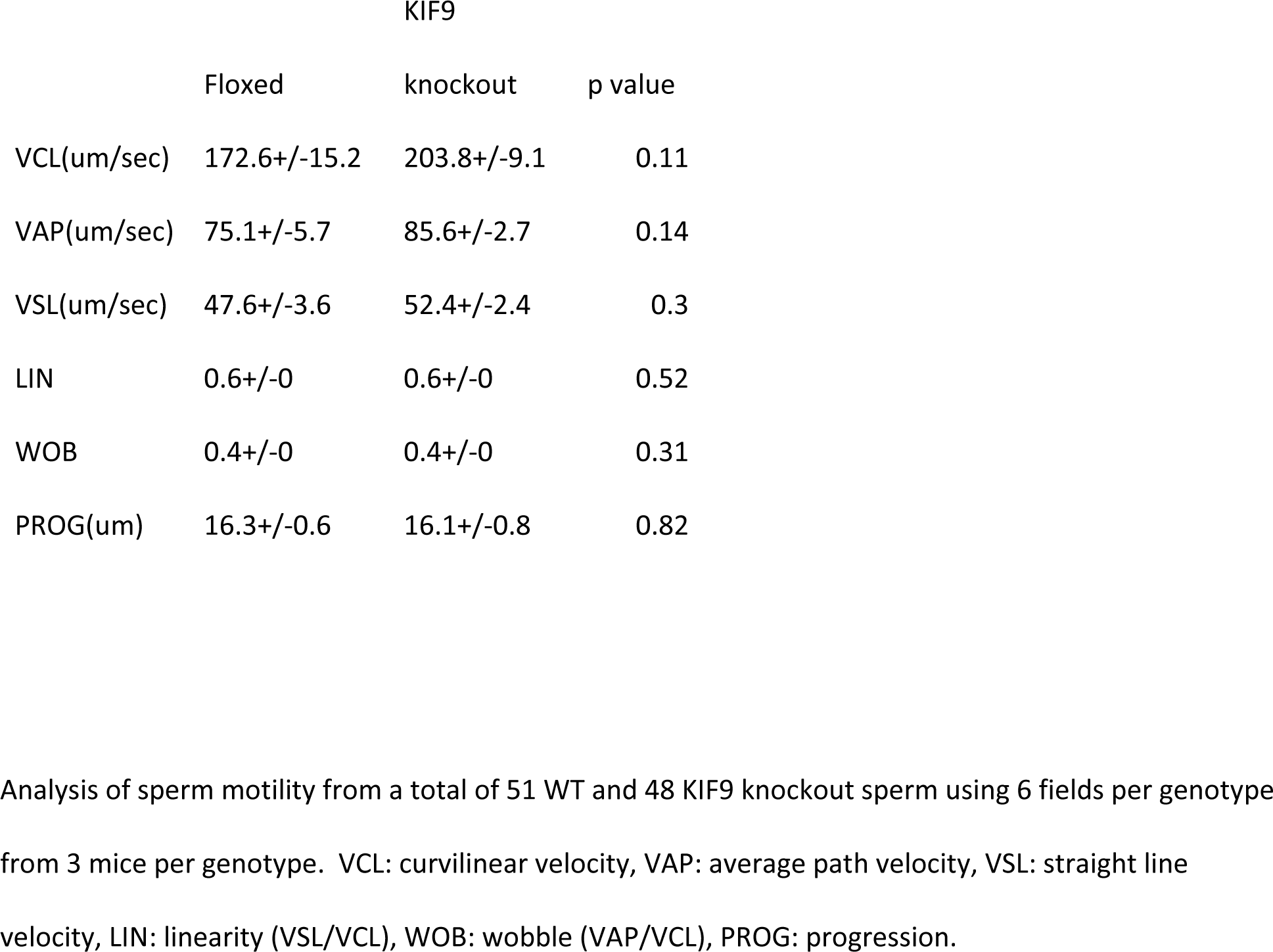
CASA motility parameters.

**Figure 1.**
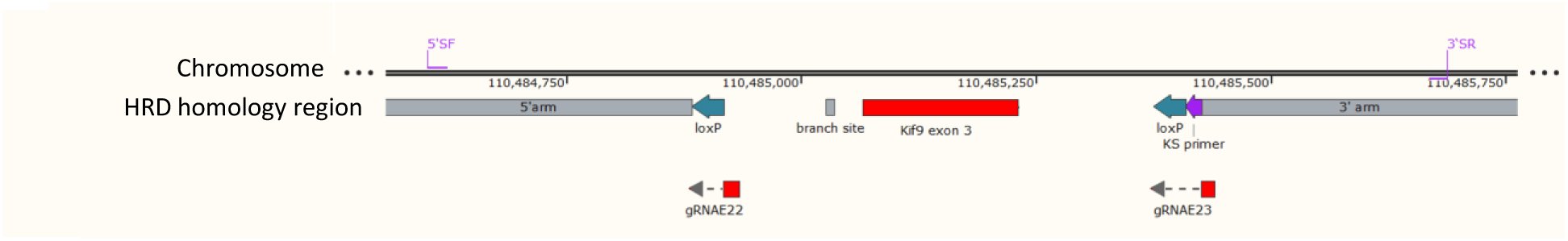
Map of the 1 KB region surrounding exon 3 of the mouse KIF9 gene. The 5’SF and 3’SR primers are used for genotyping. The HRD homology region shows the 5’ and 3’ arms used for homologous recombination along with the inserted loxP sites. gRNAs 22 and 23 were used with Cas9 to induce the recombination.

### In vitro fertilization assay

Oocyte donors were superovulated with pregnant mare serum (PMS) and human chorionic gonadotropin (HCG) for 72 and 24 hours (respectively) prior to date of IVF. On the procedure date, the males were dissected and sperm collected in HTF dish (preincubated at 37 °C, 5% CO2). Fourteen hours post-HCG injection, oocyte masses were collected in IVF HTF dishes (preincubated 37 °C, 5% CO2). Sperm were added after a one hour pre-incubation to each IVF HTF dish. After 4 hours incubation, fertilized oocytes were washed and cultured overnight at 37 °C, 5% CO2. The next morning, embryos were counted and evaluated. The fertilization rate was determined by calculating the total number of two-cell stage embryos divided by the total number of oocytes obtained.

### In situ hybridization

KIF9 mRNA was detected using the Advanced Cell Diagnostics (Newark, CA) BaseScope Red Detection kit (322971) and a 2ZZ BaseScope probe for KIF9 (710621, BA-Mm-Kif9-2EJ) targeting NM_010628.3, bases 214-402. The protocol followed the manufacturer’s instructions. In brief, paraffin tissue sections were deparaffinized, treated with hydrogen peroxide, followed by target retrieval using RNAscope target retrieval agents (322000) and protease III, and probe hybridization at 40 degrees C for 2 hours and then washed, amplified and detected using the BaseScope Red detection reagents, and counterstained with hematoxylin.

### KIF9 antibody generation and immunohistochemistry

An anti-KIF9 rabbit polyclonal antibody was generated using Pierce Custom Antibody Services. The peptide RVRPTDDFAHEMIKYGED was synthesized, conjugated to KLH and used to immunize 2 rabbits (2137 and 2138) with boosts at 14,42 and 56 days. Production bleeds from 75 days were purified using the peptide and used for immunohistochemistry. Immunohistochemistry was performed by the Histology and Comparative Pathology core facility. Paraffin sections were deparaffinized, antigen retrieval performed and then incubation with various dilutions of antibody followed by detection performed. Comparison between WT and KIF9 knockout testes led to the choice of using the affinity purified antibody from rabbit 2137 at dilutions of 1:2500 to 1:5000 dilution.

### Sperm motility analysis

Sperm were harvested from cauda epididymis under prewarmed light mineral oil (ES-005-C, Millipore) and incubated in L15+20 mg/ml BSA (Life Technologies 21083027 and Fisher BP1605) for 30 minutes at 37 deg. The sperm were diluted in L15/BSA and inserted into a MicroCell (Vitrolife 15424) and time lapse movies taken at 37 degrees at 60 fps using a 20x objective and green filter with an iphone 6plus and IDu Optics LabCam adapter. Motility parameters were measured by manual frame by frame tracking of the sperm heads and analyzing their motion in ImageJ using a modification of the CASA plugin developed by Wilson-Leedy and Ingermann[8]. For following the straightness of sperm tails, the tail morphology of the motile sperm used for the CASA analysis was traced in the first frame of each video. The xy coordinates were used to calculate the straightness ratio: the net distance from the head 10 um along the length of the tail divided by 10 um.

### Functional Enrichment Analysis

RSEM expected_count (DESeq2 standardized) dataset from 165 normal testis samples was obtained from UCSC cohort: TCGA TARGET GTEx (https://xenabrowser.net/datapages/?cohort=TCGA%20TARGET%20GTEx&removeHub=https%3A%2F%2Fxena.treehouse.gi.ucsc.edu%3A443) for gene-expression analysis. Gene set enrichment analysis (GSEA) was performed to examine KIF9 related enriched pathways in Hallmark gene sets, Reactome gene set, GO gene sets including biological process, cellular component and molecular function. GSEA was performed between two groups exhibiting high (top 10%) and low (bottom 10%) levels of KIF9 expression. Relationships were considered significant at P < 0.05 and a false discovery rate < 0.1.

## Results

To develop a KIF9 knockout mouse, a floxed exon 3 HRD plasmid was designed and inserted into fertilized C57BL6/J eggs using CRISPR gRNAs (Figure 1). Founder mice with the appropriate floxed exon 3 insertion were identified by pcr and maintained on a C57BL6/J background. To develop a whole animal knockout mouse, the floxed line was bred with C57BL/6 mice carrying cre driven by the Zp3 promoter and pups with a deleted exon 3 allele were identified by pcr. The mice were then bred further to produce homozygous whole animal KIF9 knockout mice. Both male and female homozygous KIF9 knockout mice were generated at normal frequencies from heterozygous parents and grew without dramatic differences from heterozygous littermates.

However, it was not possible to develop a pure KIF9 knockout colony because the efficiency of breeding homozygous knockout males with homozygous knockout females was extremely low. To evaluate this further, we quantitated the mating efficiency for various combinations (Figure 2A). Heterozygotes mated at frequencies and with litter sizes similar to floxed founders. Also mating of homozygous knockout females to either heterozygous knockout or floxed males was efficient. However, mating of homozygous knockout males to either heterozygous knockout or floxed females was produced very few pups. One-way ANOVA indicated a highly significant difference between groups (p<.0001) and T-tests indicated significant (p<.0002) differences between M-- matings and either F-- or M+-/F+- matings. To determine if the mating defect was due to a reduction in the ability of sperm to fertilize eggs, an in vitro fertilization assay was performed (Figure 2B). Sperm were isolated from knockout males with similar yields to wild-type and were motile. However, the KIF9 knockout sperm were strongly defective in their ability to fertilize eggs (p<.02).

**Figure 2.**
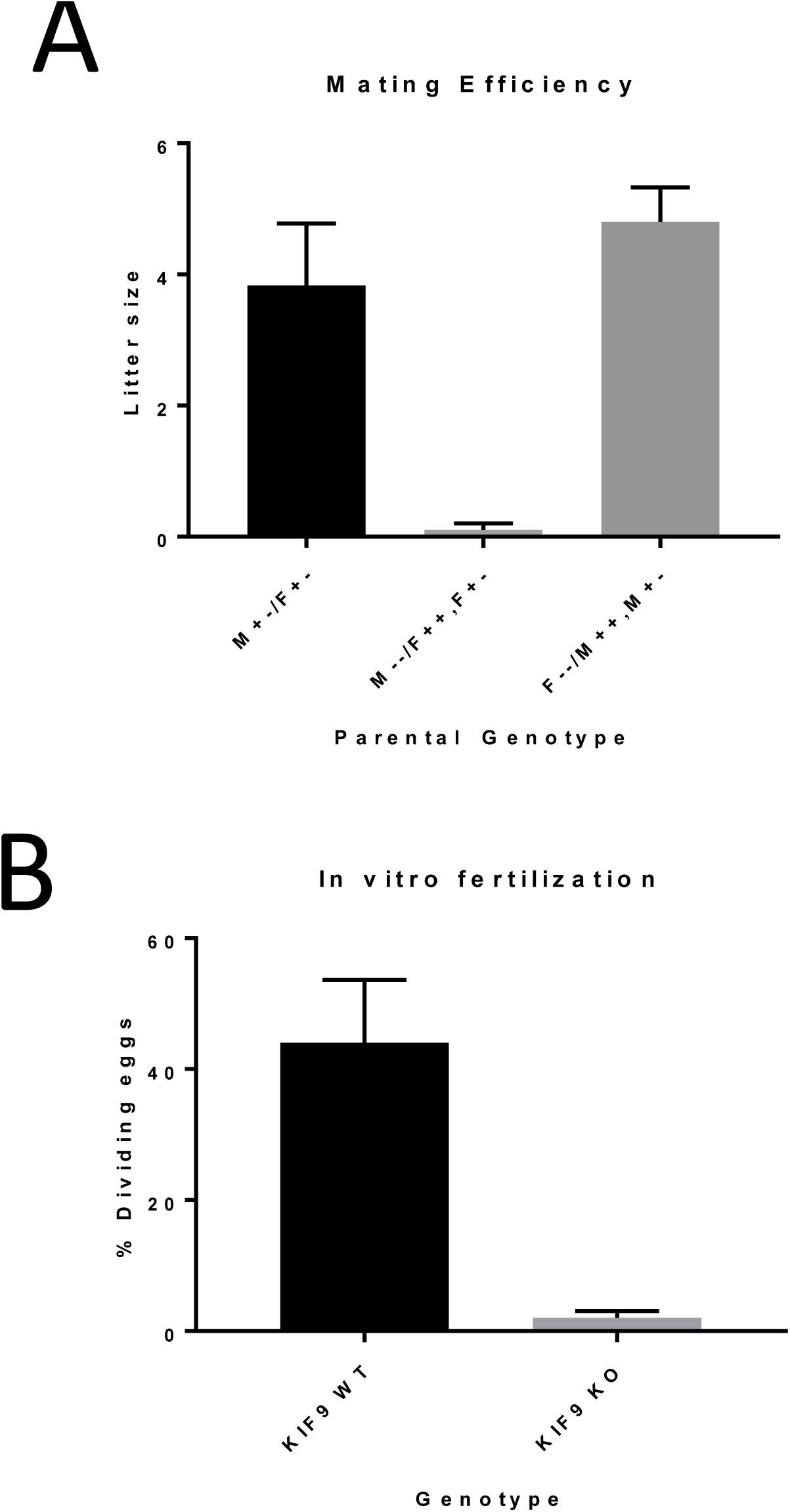
A. Homozygous male KIF9 knockout mice (M--) are defective in mating. To test the mating efficiency of various genotypes, comparisons were made between crosses of males heterozygous for KIF9 (M+-) mated with heterozygous females (F+-) (left column), homozygous knockout males (M--) mated with either wild-type (F++) or homozygous (F--) females (center) or homozygous knockout females (F--) mated with either wild-type (M++) or heterozygous (M+-) males. Data are means and SEM of 6, 10 and 15 matings for M+-/F+-, M--, and F—genotypes, respectively. One-way ANOVA indicates a highly significant difference between groups (p<.0001). T-tests indicate significant (p<.0002) differences between M-- matings and either F-- or M+-/F+- matings. B. In vitro fertilization efficiency of sperm from wild-type (KIF9 WT) or knockout (KIF9 knockout) males were compared. Data are means and SEM of tests of sperm from 3 mice of each genotype, and the means are different with p<.02.

We then examined the expression of KIF9. The Human Protein Atlas, GTEx and FANTOM5 datasets all indicate that KIF9 mRNA is most highly expressed in the testis both in human and mouse (Figure 3A). We then performed in situ hybridization of a number of mouse tissues, and found indeed that KIF9 mRNA is most highly expressed in spermatocytes of the testis (Figure 3B). Interestingly, expression of KIF9 mRNA was also found in specific regions that have multi-ciliated cells such as the ependyma, bronchioles and oviduct. Comparison of KIF9 knockout testis with wild-type revealed that the mRNA expression seen in the seminiferous tubules was lost in the KIF9 knockout (Figure 4A). To evaluate protein expression levels, an antipeptide antibody to KIF9 was made against the sequence RVRPTDDFAHEMIKYGED which is in exon 2 upstream of the deleted exon 3. Staining of the seminiferous tubules with the antibody confirmed that KIF9 was expressed in developing spermatocytes in the WT and lost in the KIF9 knockout spermatids (Figure 4B). The level of signal in sperm was too low to determine whether KIF9 is present in sperm tails or not.

**Figure 3.**
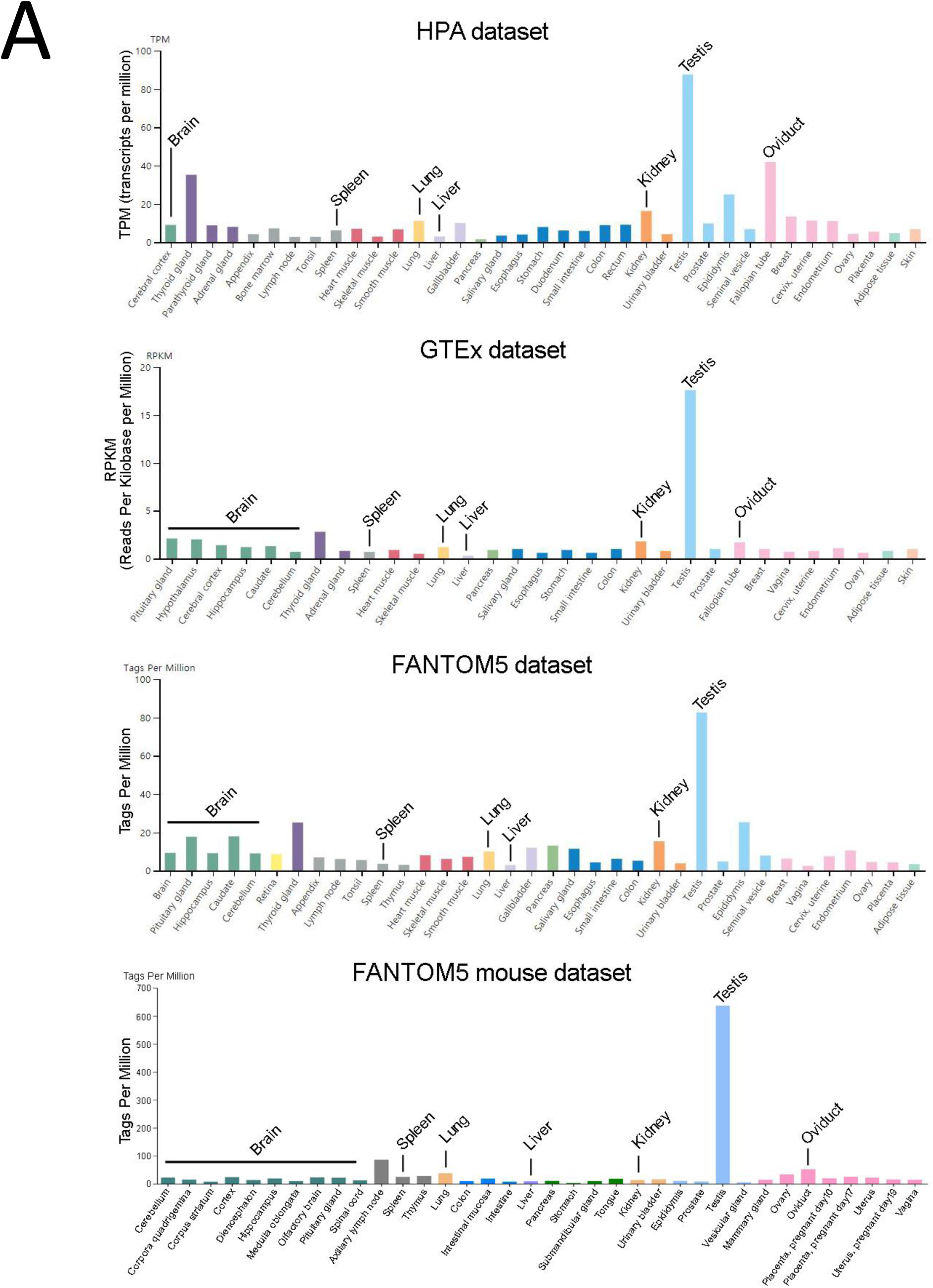
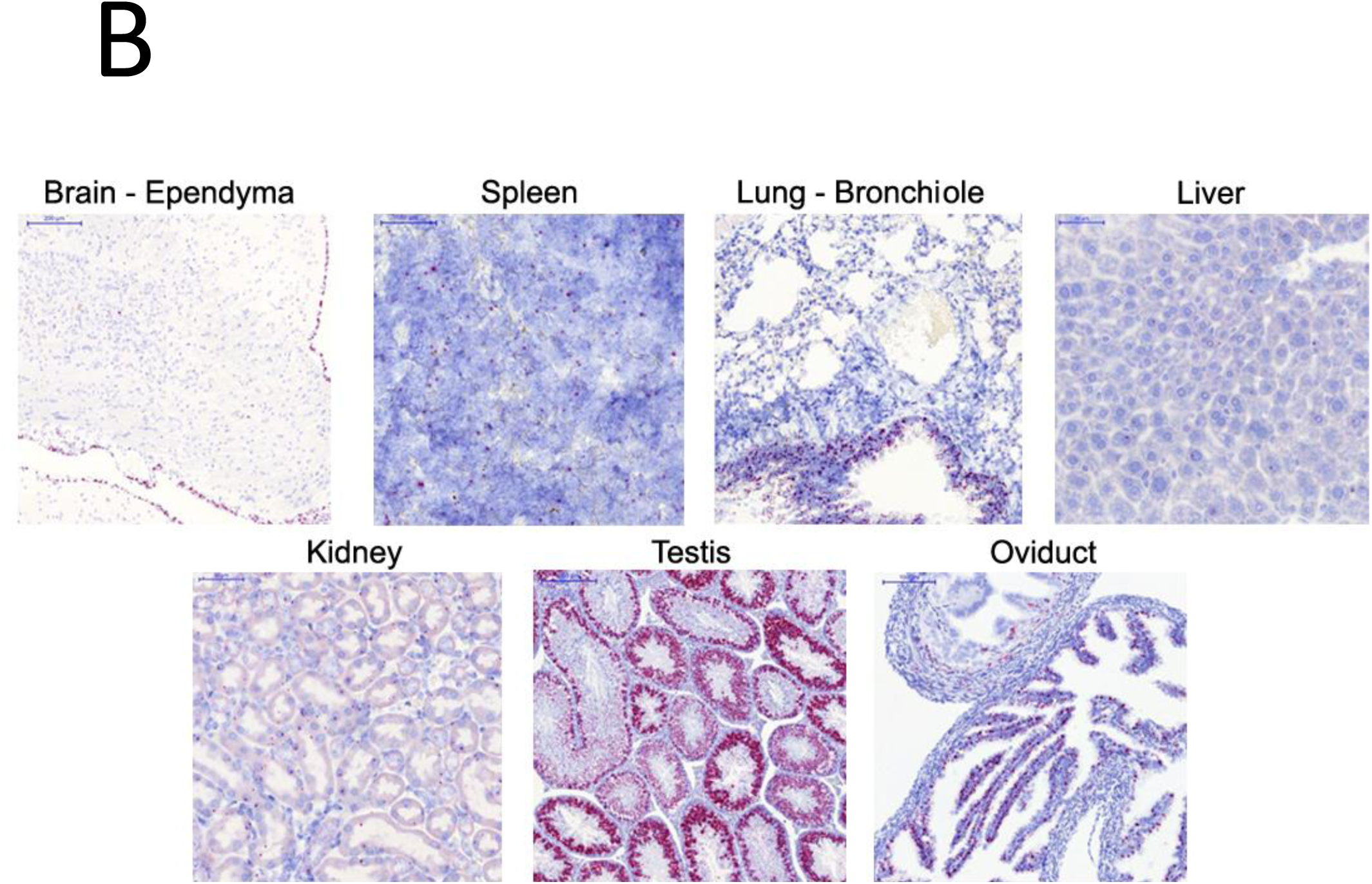
Expression of KIF9 mRNA in normal tissues. A. mRNA expression of KIF9 in organ tissues reported in The Human Protein Atlas (HPA) dataset (http://www.proteinatlas.org), the Genotype-Tissue Expression (GTEx) dataset (https://www.gtexportal.org) and Functional ANnoTation Of the Mammalian genome5 (FANTOM5) dataset (http://fantom.gsc.riken.jp). B. In situ hybridization for KIF9 mRNA in selected mouse tissues.

**Figure 4.**
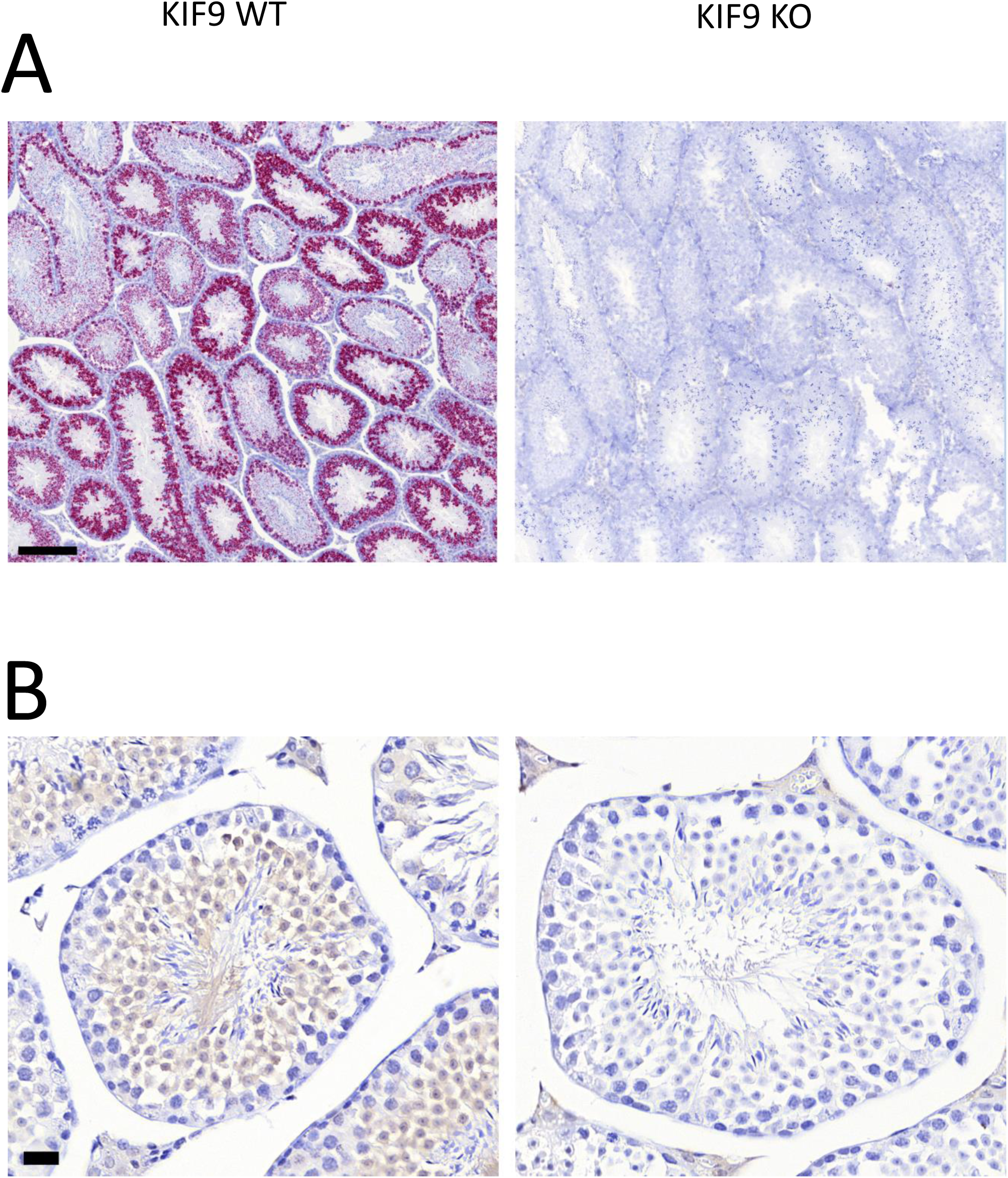
Confirmation of loss of KIF9 expression in KIF9 knockout testis. A. In situ hybridization for KIF9 in testis for WT (left) and KIF9 knockout (right). Scale bar is 200 um. B. Immunohistochemical staining for KIF9 in WT (left) and KIF9 knockout (right) testis using antibody 2138 at 1:5000. Scale bar is 20 um.

Given that KIF9 is expressed in the testis, and that KIF9 knockout sperm showed a defect in fertilization, we examining the motility of WT and KIF9 knockout sperm in more detail (Videos 1 and 2). CASA analysis showed no difference in standard motility parameters (Table 1). However, the KIF9 knockout sperm did show a subtle difference in motility. The curvature of the sperm tail appeared to be reduced in the KIF9 knockout (Figure 5A). Quantitation of the shape of the tail confirmed an increase in straightness of the tail in the KIF9 knockout (Figure 5B, p = .0003).

**Figure 5.**
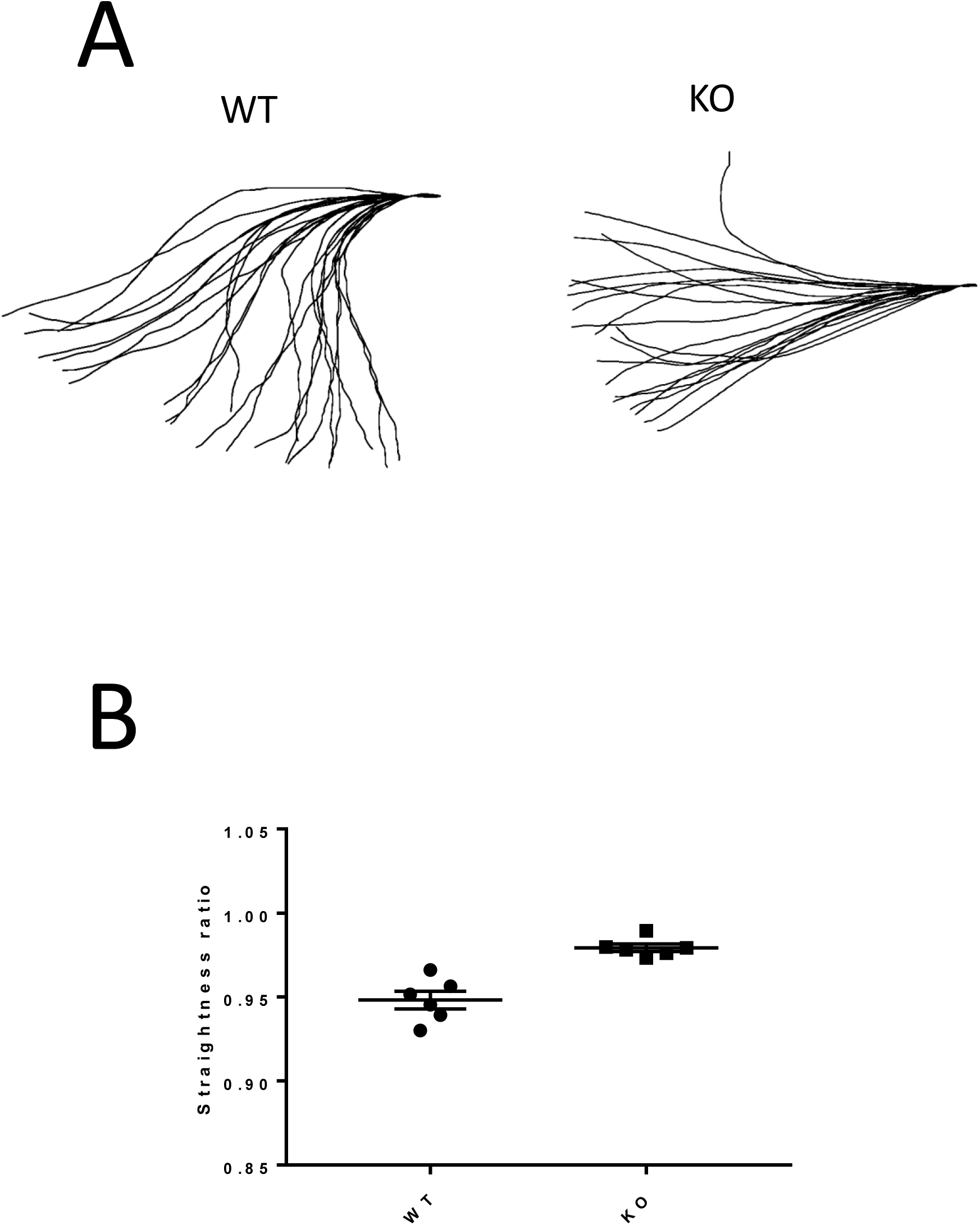
Evaluation of sperm tail waveform. A. Waveform patterns of the sperm followed in Videos 1 and 2. B. Evaluation of straightness of flagellar waveforms of sperm used for the CASA analysis. A total of 51 WT and 48 knockout sperm tails from 3 mice of each genotype were analyzed. P<.0004 (unpaired t test).

GSEA was performed to identify significantly enriched pathways that differed between the high (top 10%) and low (bottom 10%) KIF9 mRNA expressing testes based on pathways provided in Hallmark gene sets, Reactome gene set, Gene Ontology (GO) gene sets including biological process, cellular component and molecular function (Supplementary Figure 1). A number of significant correlations were identified, including fertility, cilium assembly, cilium movement, and ATP-dependent microtubule activity. These correlations are consistent with a role for KIF9 in sperm motility.

## Conclusions

A KIF9 knockout mouse was generated through deletion of exon 3 of the KIF9 gene. Knockout mice were viable and generated at frequencies similar to wild-type from heterozygous parents. However, breeding of homozygous knockouts revealed that male knockouts showed a breeding defect while female knockouts did not. In vitro fertilization assays revealed that the sperm from KIF9 knockout males were strongly reduced in fertilization efficiency. However, most motility parameters of the knockout sperm appeared normal, with the major difference that was observed being a change in the waveform pattern. Consistent with the mating phenotype, expression of KIF9 was high in the testes in wild-type mice and lost in the knockout. Other cells that express KIF9 at the mRNA level include lining cells of the oviduct, bronchioles, and ependyma.

Phylogenetics studies reveal that KIF9 is present in species that form cilia or flagella[9], consistent with our observations that KIF9 is highly expressed in the testis as well as detected in other regions that generate cilia. KIF9’s homologue in Chlamydomonas reinhartii, KLP1 [5], is important for flagellar motility. Knockdown of KLP1 in this organism leads to slower or immotile flagella [6]. Knockdown of the KIF9 homologue in trypanosomes [10] or fern gametophytes [11] results in reduced motility.

At the mRNA level, FANTOM and HPA databases indicate the highest KIF9 expression is in testis. Our in situ hybridization analysis confirms high levels of KIF9 mRNA in testis particularly in the developing spermatocytes. We confirmed the presence of KIF9 at the protein level in the spermatocytes using an antipeptide antibody. Proteomic studies have revealed the presence of KIF9 in human sperm[12-14] as well as sperm of several mouse species[15]. Thus, based on studies in lower organisms and localization in mammalian sperm, we anticipated that there would be a strong motility defect. Although we found that KIF9 knockout sperm had a greatly reduced ability to fertilize eggs, the motility of the sperm in vitro is not strongly inhibited as measured by CASA analysis. A subtle alteration in waveform was observed, with the knockout showing a straighter waveform. Our current data cannot resolve whether the defect in fertilization reflects this altered swimming pattern or possibly other structural changes in the sperm. It is interesting to note that a more rigid swimming pattern was also noted for sperm lacking the sperm specific isoform of calcineurin[16], along with a defect in fertilization capability. KIF9 has been found to be phosphorylated[13], and thus it is possible that regulation of KIF9 by phosphorylation contributes to the regulation of the flagellar waveform.

Our GSEA analysis in testes was consistent with the above findings, showing that high KIF9 has significant correlation with sperm motility. In addition, high KIF9 was also related to spermatogenesis, which may explain the high staining of KIF9 in developing spermatocytes and the loss of fertility in the knockout male mouse. High KIF9 showed also a significant association with cilium in biological process and cellular component, supporting a possible direct role of KIF9 in ciliary motility.

Another study also recently reported a male sterility defect in KIF9 KO mice with the sperm showing an altered waveform [7]. Interestingly, in those mice although the mating defect seemed weaker, a stronger motility defect was observed, including significant reductions in path velocity, straight line velocity and curvilinear velocity. The differences in results could be due to different knockout strategies.

Our in situ hybridization analysis confirmed that other tissues showed expression of KIF9 at the mRNA level, consistent with the FANTOM and HPA databases. Interestingly, in a number of cases, the expression was in epithelia that contain multi-ciliated cells, such as bronchioles, ependyma and lining epithelium of the oviduct. The functional consequences of KIF9 expression at these sites is unclear. Female KIF9 knockout mice showed normal fertility, and the knockout mice did not show indications of greater sickness, or shorter lifespan.

KIF9 mRNA was also evident at low levels in areas of kidney, spleen and liver (hepatocytes) that are not known to contain multiciliated cells. These tissues can contain primary cilia, however, and the low levels of KIF9 could reflect a potential function in primary cilia. In addition, KIF9 has been reported to be important for matrix degradation in macrophages[1] as well as interacting with GEM with possible functions in spindle control[2, 17]. The low levels of signal detected in tissues that do not have multiciliated cells may reflect such alternative functions.

GWAS studies up until now have not identified a consistent association for KIF9 with a particular disease. Isolated reports suggest possible associations with Huntington’s disease[18] and breast cancer[19]. Interestingly, increased KIF9 expression has been correlated with shorter survival time in glioblastoma including increased EMT and immune function [3]. Further studies will be needed to clarify the function of KIF9 in these pathologies.

In summary, we have generated a knockout mouse lacking the kinesin family member KIF9. Whole animal knockouts are viable with the strongest phenotype being male sterility. Additional more subtle phenotypes may be identified, given the expression of KIF9 in tissues containing multiciliated cells as well as other tissues. Given the lack of phenotype other than male sterility, inhibition of KIF9 function could be considered as a possible approach for a male contraceptive.

## Supporting information

Video 1 KIF9 WT

Video 2 KIF9 KO

## Acknowledgements

We thank Hong Zhang for assisting with immunohistochemistry and Kelvin Davies for comments. We acknowledge the use of the Cancer Center Core facilities, including the Analytical Imaging Facility, Genomics, Histopathology, and Animal facilities. JES is the Betty and Sheldon Feinberg Senior Faculty Scholar in Cancer Research at the Albert Einstein College of Medicine and is a member of the Gruss Lipper Biophotonics Center.

## Author contributions

YZ generated the KIF9 floxed mice, KC performed the in vitro fertilization assay, SYC performed GSEA and database analyses, AB performed pathological analysis, and JES performed breeding and motility studies.

