## Supplementary figures and images for "A role for KIF9 in male fertility"

### Supplementary Figure 1

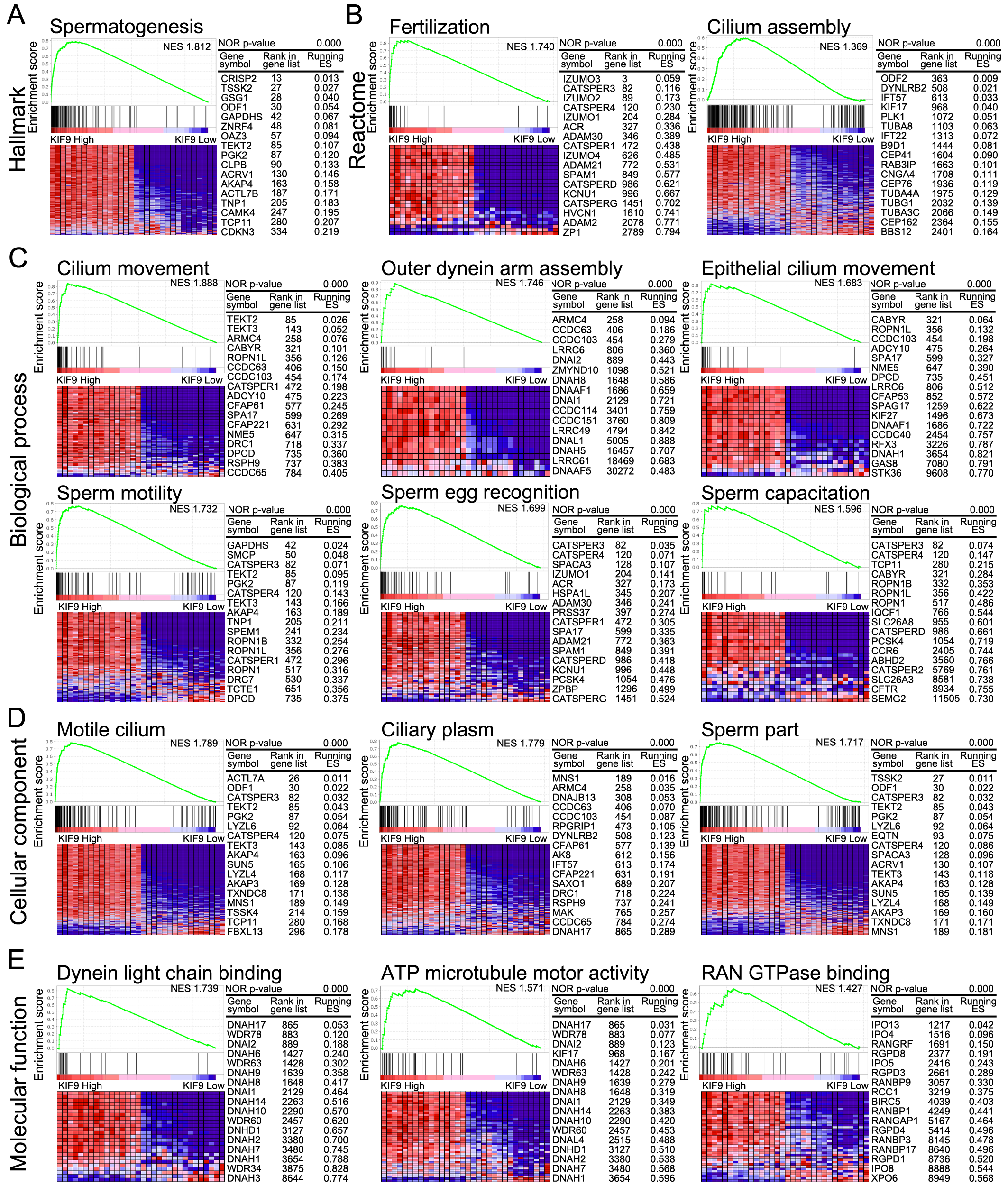
