## Supplementary Information for "A role for KIF9 in male fertility"

**Supplementary Data**

Supplementary Figure 1 Legend: GSEA based on hallmark gene sets, Reactome gene sets and GO gene sets between KIF9 High and Low groups. A. Representative GSEA data with p value and normalized enrichment score (NES) in Hallmark gene sets. B. Representative GSEA data with p value and NES in Reactome gene sets. C-E. Representative GSEA data with p value and NES in GO biological process, cellular component and molecular function gene sets respectively.

Video 1: KIF9 WT sperm. Bar in first frame: 20 um. Frame rate slowed from 60 fps to 10 fps.

Video 2: KIF9 knockout sperm. Bar in first frame: 20 um. Arrow indicates sperm tail traced in Figure 5A. Frame rate slowed from 60 fps to 10 fps.

Supplementary Table 1: primer sequences

| | |
|---|---|
| gRNAE22 | agagagttggagttatgttt agg |
| gRNAE23 | atgttctaccactagatcac agg |
| 3'SR primer | gatatgcaaagggtgggttg |
| 5'SF primer | tgcatacacattcgtgaaaaca |
| KS primer | cgataccgtcgacctcg |

Supplementary data file 1: HRD sequence

tcgtggtggaaaaatcacaaagcagctgaaagacaggaagcacgtgtacttacagggcagggactgtgggttgtgattcatctcttgtagccaaggct
gtcctccaactcacaatggagctgaagctggccttgaactccaatccagctgcctttacctcccaagtattagggttgcaggtgtgcaccaccatgctag
gcttttattctgatcttattgaatggcatctccctcctggttggtatttgatgatggctactaaaaatattgtcgttctcctttctccagctaaaatgggtact
aggaaaaaggttcaagcgtttgttcgagtcaggcctactgatgactttgctcatgaaatgatcaaatatggagaagacaacaaagtaagtgcagtgtg
tttgactttgtttatgcagaccttcccttttcacacagccatttaaagaccgccacagagggactggagcgagagctcggtgggtacagatgccggcca
gcatgcagcatgcctgagggcctgaggactgtcccctccatccgaatggtggagggaaggaaccaactcctaagagttgtcttccaaattccatacac
atgctattgtgtgcacatttacacacacacacacacacacacacacacacaattttcatcaaaggggctaagacagaattgatagatgttcatgta
aactctagagtcagtccatcaactccaaactctgctgttagtgaggaatttcttgagattctcattttttttaatgtgttcatatggcgggaggagcacact
ggtgtgctccagctcaggtgtggaagtcagagagcagcttgcagaactgtagtttctcttctctcatgacgtgggtgctgaggaatcaaacttggattgt
ttggttggtagcaggcacctttcttcactgaatcatcttgctggccctcaggtattttaaatgaatagctgaagaataaattgctcaggtgctgaaagttc
tggtagtcataaaagagctcagggatggccctgtgtaatgtgctctatggcatgatttaaacaaacatggacccacagagcatgaaattttctcagca
gcacttgctatggggtattcatttgtttggattttgttggagtcggggtctcatgtaacccagctggctttgaactggctgtttaaaatagggtaactatgg
cccttgtgcttttacttcccaagtgctgggattgtacctacaaatcaccctgataaccctatctgtcttagccttagttttgtttttctatggacagctaggtc
ttggttcttctattacagtagccagtagaagatttcagggaaagccccatcaggagcttggggaataagctggatatatatggtgctgtaccatcatctg
aacagttgggagactgagaggaaaatccaagtacctaaattcaaagcgagccttcatatcacaaaagtgacatttaaaaaaggcttggagtctcact
gggaaatggctggaggtgcatttgaaattctgctggtcacaagtaggttaggccccctccatttacattaacataagtgactacttgtatctgaacactt
gatcatcccaccattgctcccagtggctcccagctgaggaagtaagcgctggccatagcgcagaggatgcagaggcatcagtcacacagacagggcc
tgcccgggccgttgtttccagtttccctgttgctttgcctaagtcaagactcagtagttaaaagcaccgggtgctcttccacaggacctgtgcttagtaca
cagtacccacaaccatacatacatatactacatgcatacacattcgtgaaaacatgttaaaaaacaaaaaagtaaatttaaaaactttttttaagcctc
acatcctctgttatttctaagttggactttactgaccagtgattgattatttatttgtttgtttgtttgtttgtttgtttgatacagtctccctatgtagccctag
ctgtcctggaatgttctatgtaaatcagactgcccttgaactcacggagatccacctgcatctgtctcctgagtgctatcagtgattccttttacaaaggc
ttcctaaaATAACTTCGTATAATGTATGCTATACGAAGTTATcataactccaactctctcttatttttgtgacatttcacacatgttcccca
caatggcctgaaactccagagcctcctaacatagctgtccggggctgggttcaagtgtgtgtcacccacacctgttgtctatctctctctcccccctaga
gcatcgacatccacttgaagaaggacactcggagaggggttgtcaataaccagcagacagactggtccttcaagctggatggagtgcttcacaatgc
ctcccaggacttggtttatgaaacagttgccaaggatgcggtttctcaagcccttgatgggtacaatggtaatttaattaactgttactttatctaggact
ctgaggatccccacttggaggaggaggtacaagatgtactctatcctgaataaaacaacacaaaagaacttcatttacttacattgtaaacttacttgt
cagcctgtgATAACTTCGTATAATGTATGCTATACGAAGTTATCGATACCGTCGACCTCGtagtggtagaacatagatttactta
atgacattgtaggcatggcttcatggcttcatcagtctctgagatgttacctttattggattgttagacgatcttggctttttttatcattaattaattagttta
ttcactttacatcctgattgaagcctctctatcccccccccgtcctcctagtcccaccctcgcacccctacccccattccccatcacagaaggggagaccc
cccaatagtaccaacccaccctttgcatatcaagtcacagcaggactaagtgcatcctttaccactgaggccagacaaggcagcccaactacgggaa
aaggacccaaaggcaggcaccatagtcagagacagccccactccaattgttagggaccctacatgaagtccaactgcacatctgctacagatgctctt
tatttggtggctcagtctctgtgagccccatggacccaggttggttgactctctaggtcttcttgtaatgtccttgacctctccaattcactcagtcgttctc
ccaactcttccaaaggaatccctgagctccacctattgtttgggtgtgagtctctgcatctgcttctgtcagctgctggatcaagcctctcagatggcagt
tatgccaggctcctgtctgcaagcatagcagagtatcattcatagtgtcaggggttggctctctcccatggggtgaagactctggcttttcatgggtcttc
aatggtcaactttaggcttgtttcaagcctctcaagcagtgtgtgtgtgtgtgtgtgtgtgtgtgtgtgtgtgtgtgactgtgtctatccgaatctcagtctc
actatgtagccctgtctggcctagaactcactatgtagacccagctagccttaaactctcagaactctgcctgccaccactgcctggatttttttttttaag
attaaattcttgtttgtttgtttggtagtgttttgttttgtttttttagacagtatttcactgtgtagttctggctgtcctggaacttgctctatagaccagatctg
gaactcatgaagatcagcctacctctgcctcccaagttctgggataaagggtgtgtgccaccccacttagccctacatgtgatttttttttttttaggtatt
tatttcatttacatttccagtgctatcccaaaagtcccccacatgctacctacatgtgatttttaaaaacactttaaaagtagatgggactggagaaatg
actcatcagttaagagcactggctgcatgctcttccaggggacccaggttcaattcctagcacccacaatgcagctcacaaccatctataactccagttt
caaggggaccctcttctgtccaatttggacacccacctggcactcacatgatccacagacatacgtgcaggcaaaaccccatgcgtttaaacattactt
taaagcagaacaaaccccagcacccttgtgtgtggcatagtagcaaattcctataatttcagtgtttagaagcagaggcaggaggatgaggagtgta

aagtcatcctcagctacctaccagttagatgtcatcttgagctacatgtggcactatctaaaaataaataaagcttaaaatataaaactaaagttgcag
tttgaagactgccgtctggtggctgctgatcggaatgttctcagtttgttggttagttgcttggtactgggtttgaaacccagagtctcacaaatattaagt
gcatgttctctcaccacactatatgtgtagctccaaaagttttagactcctctgtaggaaagcccatggagggagccaggcgtggctgtgcatgcttgta
catccagcacttgggaggtgacgcaggatgaccagggatttgagaacagtttggattatatagctagacttggtttcaggaaaatccagtgactcttga
gttcccaaaattgtaatgtattcttcatgtgatacataggtgtggttcccactgacccaatgatatattcatgattccctgagaggctctgtggcctggag
ataaagatggaggggg
